# Inferring context-specific site variation with evotuned protein language models

**DOI:** 10.1101/2025.02.19.639211

**Authors:** Spyros Lytras, Adam Strange, Jumpei Ito, Kei Sato

## Abstract

Multiple sequence alignments (MSAs) have been traditionally used for making inferences about site-specific diversity in proteins. Recent advancements in the field of artificial intelligence have highlighted the potential of protein language models (pLMs) to capture similar protein properties. Unlike MSAs, pLMs can make inferences from single sequences, without the need for a set of aligned sequences. In this study, we introduce a novel, pLM-based metric, termed ‘pLM entropy’, to assess protein site conservation and variability. We test this metric using versions of two popular pLMs (ESM-2 and protT5) fine-tuned on the diversity of different Influenza A virus serotype hemagglutinin proteins. Our study demonstrates how our pLM entropy metric can capture which sites are more likely to change in a specific sequence context and how fine-tuning pLMs on a set of evolutionarily related proteins (evotuning) can improve the models’ understanding of the group’s diversity.

## Introduction

The sequence diversity between evolutionarily related proteins reflects their functional properties and constraints. Aside from neutral evolution, protein sites that do not change throughout evolutionary time have a conserved role, while sites that change frequently can be under diversifying selection (1). Traditionally, this diversity in protein sites has been studied by inferring multiple sequence alignments (MSA), a hypothesis of sequence homology between sequences that once shared a common ancestor. One common quantitative measure to assess site-specific conservation or diversity in protein sequences is calculating the Shannon entropy of the frequency of each amino acid at a given site in the MSA (2). In this metric, lower values indicate higher conservation, while higher values reflect greater diversity (i.e., higher mutability). Since entropy values are derived from an MSA, they represent the average effect across the sequences in the alignment. However, the strength of evolutionary constraints at a given site is inherently context-specific and may vary among sequences within the MSA. If we can determine entropy values associated with each individual sequence, then we could compare the strength of conservation or variability for a specific site across different sequences, enabling the detection of context-specific evolutionary constraints (3).

Recent advances in the field of natural language processing (NLP) have become increasingly relevant to studying biological sequences. The same way that grammar and syntax determine how words are arranged in a sentence, evolutionary relatedness, and functional and structural constraints condition how amino acids are arranged in proteins (4, 5). This similarity in their context-dependent behaviour constitutes deep learning model architectures such as Transformers—originally developed for NLP applications—also well-fitted for studying interactions between protein sites (6, 7). Consequently, the concept of large language models (LLMs) has been borrowed from the NLP community to apply on biological sequences, leading to the development of many protein language models (pLMs) (8–13). The majority of popular pLMs developed so far leverage a wide diversity of known proteins for their training datasets (primarily retrieved from the UniRef database (14)) which allows them to retain biological features associated with sequence patterns (15, 16). The pLMs’ ability to relate sequence context to function and structure in such a multidimensional representation space enables powerful applications, from efficiently predicting 3D protein structures (10, 17, 18) to inferring fitness effects of amino acid substitutions without the need for a sequence alignment (19).

The probabilistic nature of pLMs allows them to make site-specific inferences by considering the amino acid context across all the sequence’s sites. This includes inferring the likelihood of each amino acid at a given site in a protein sequence. Hie *et al.* (2021) (4) applied this pLM property to viral proteins, referring to the inferred per site probabilities of having any amino acid as a measure of “grammaticality” (as a reference to natural language). So far, this metric has mostly been used for inferring the likely phenotypic effect of specific amino acid substitutions (20). Here, we extend this concept by introducing “pLM entropy”, a novel metric that represents context-specific sequence site variability and conservation. While pLM entropy correlates with traditional MSA-based entropy, it uniquely captures context-specific evolutionary potential at each site without having to rely on a sequence alignment.

The extent to which a pLM can understand the context of a given protein depends on how well represented that protein group’s diversity is in the model’s training dataset. Most popular pLMs are trained on a wide diversity of primarily non-viral proteins. Indeed, existing pLMs-based methods for fitness inference tend to underperform specifically on viruses (23). Sequence likelihood inferred from pLMs are biased by the sequence’s phylogenetic distance to the species most prevalent in the model’s training dataset (24, 25). This means that if the vast majority of proteins a pLM is trained on are human, then the resulting model will perceive proteins from species related to humans (e.g., primates) as more likely than more distant, under-represented species. Similarly, if foundational pLM training datasets have proportionally fewer viral proteins than sequences from the rest of the tree of life, the models’ understanding of the viral diversity will be limited. Instead of training new pLMs from scratch on different datasets, a solution proposed by Alley *et al.* (2019) is to fine-tune existing models to a local evolutionary context, referred to as “evotuning” (26). This approach has been shown to improve model understanding of a given group of proteins and improve performance of predictive tasks for that group (26, 27). However, there are no clear criteria for selecting sequence datasets for evotuning, nor are there established metrics to assess the extent to which evotuning enhances the ability of pLMs to capture the context of target proteins. In this study, we demonstrate that pLM entropy can serve as a useful metric for comparing pLM performance. and provide valuable insights into the selection of sequence datasets for evotuning.

As a study protein group, we focus on the Influenza A virus (IAV) hemagglutinin (HA) protein for a number of reasons; First, IAV is an important viral pathogen that has caused four pandemics in the past century and continues to cause substantial disease and mortality both in humans and animals (28). The HA is the protein responsible for viral entry into the cell and the primary target of host adaptive immunity, hence it contains substantial functional diversity (29). As a result of the virus’s impact on global health there is a large number of diverse sequences collected from a variety of hosts and dating as far back as the 1918 IAV pandemic. Finally, IAV HAs are separated into multiple distinct phylogenetic groups, called serotypes, allowing for easy separation of training and testing datasets. Herein, we present and use HA-evotuned pLMs to infer site-specific pLM entropy values across HA proteins of key IAV serotype groups and demonstrate how this metric holds predictive signal for which sites are more likely to change in a given sequence context.

## Results

### 1. Evotuning pretrained pLMs with the IAV HA protein

To assess the effect of virus protein-specific evotuning on pretrained pLMs, we use two popular models, both originally trained on similar datasets derived from different UniRef clusters (30) and the BFD (Big Fantastic Database) (31) but having somewhat different architectures. The first model, ESM-2, is an encoder-only model trained through unsupervised masked learning with a single mask token (8). The second model, protT5, is based on the T5 NLP architecture (32) and includes both an encoder and a decoder trained in unison. The base T5 architecture allows for the use of multiple masking tokens during training, however, a single token was used for all masking in protT5’s training, consistent with ESM-2’s training approach. The encoder component of the protT5 model is available to use independently and was chosen here for a more similar comparison to ESM-2. The two model types have been shown to have comparable performance for different evolutionary inference tasks, such as predicting site conservation, making them a useful pair to compare for our study (19). To reduce computational burden while retaining performance, we chose to use the version of each model with the smallest size that still retains comparable performance to its largest version. This criterion resulted in selecting the 33-layer ESM-2 model (650 million parameters) and the 24-layer protT5-XL encoder model (1.2 billion parameters).

ESM-2 comes with a pretrained masked language modelling (MLM) head layer for inferring model probabilities of any amino acid token being present in any site of the input protein. This is not available for the protT5 encoder, so we adapted the model by adding a facsimile MLM head to it. The ESM-2 MLM head was trained along with the rest of the model layers which is not the case with our added protT5 MLM head. To avoid having random weights on the added head, we performed 2 epochs of preliminary training using a random subset of sequences from the UniRef50 cluster (used for original ESM-2 training) without altering the weights of the existing protT5 layers. Hereafter, we refer to this adapted version of protT5 as protT5MLM.

To evotune the two pLMs, we collected a comprehensive set of unique IAV HA sequences available on the NCBI Influenza Virus Database (33). We further separated the training sequences by HA serotype and compiled serotype-specific training datasets for the four serotypes of greatest concern to human and animal health (and consequently the ones with the greatest number of sequences available): H1, H3, H5, and H7. Each serotype set of sequences and the combined set were then used to perform unsupervised masked learning (34) on both models, creating the respective: H1, H3, H5, H7, and HA-all evotuned versions of each ESM-2 and protT5MLM (Figure 1A). To assess the robustness of our approach to training dataset availability, we also created a sequence set consisting of all HA sequences collected in the earliest 80% of the distribution of sequence collection dates. This training dataset was used to evotune an additional version of the models we refer to as HA-80.

**Figure 1.**
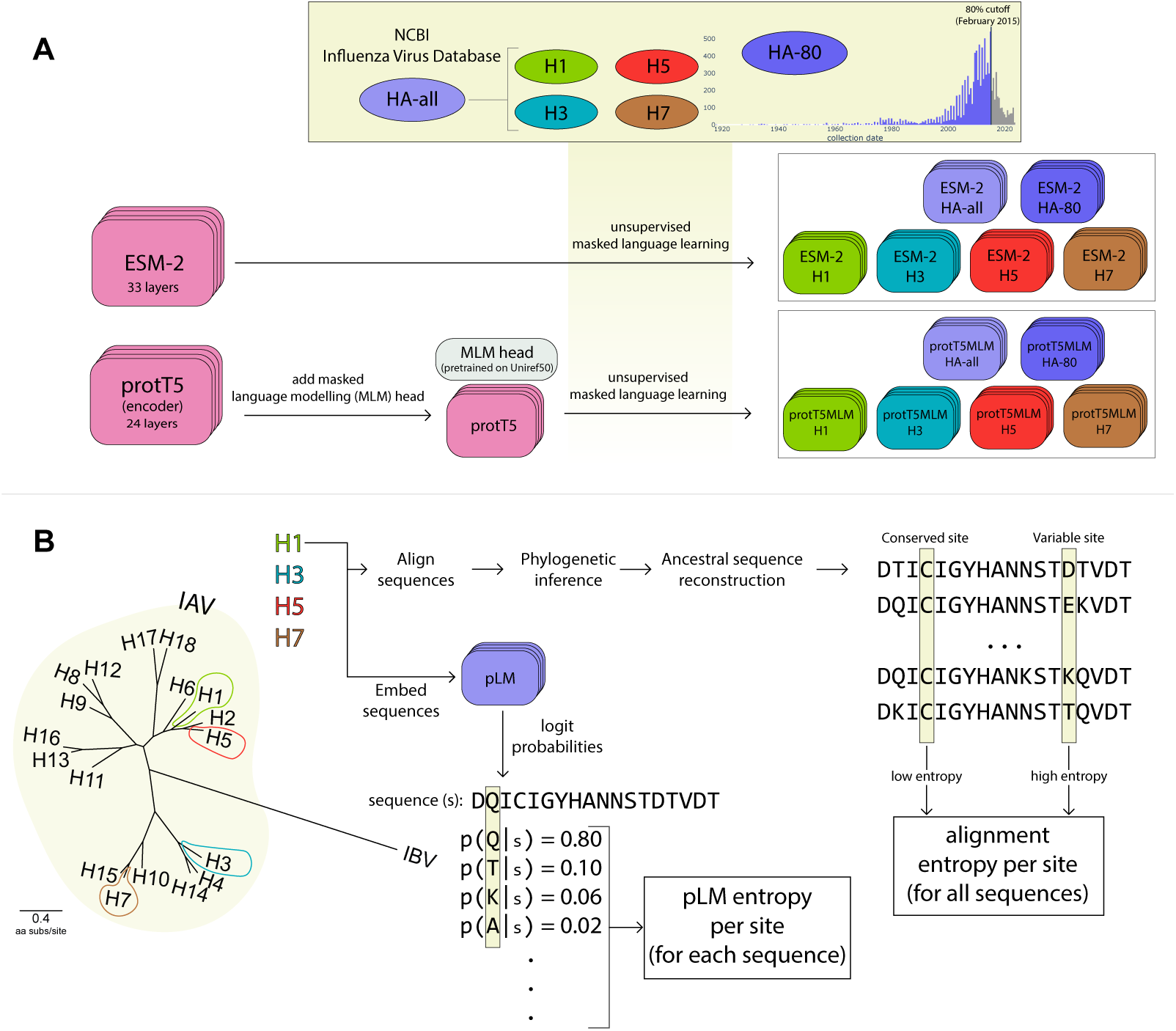
A) Schematic of training dataset and model fine-tuning strategy. B) Description of the testing dataset, for each of the H1, H3, H5 and H7 testing alignments pLM entropies per site for each sequence and alignment entropy per site for each alignment were computed.

### 2. A pLM-based metric for assessing site-specific conservation

One way to test whether evotuned pLMs have a better understanding of the diversity across the protein group they were fine-tuned on is to try to infer which protein sites are more or less likely to change. Inferring this information from an MSA traditionally involves calculating the Shannon entropy of the frequency of amino acids present in each alignment site (Equation 1) (2). Intuitively, low entropy values indicate conserved sites (an entropy of 0 meaning that all sequences have the same amino acid in that site), while high entropy values correspond to variable sites. However, this MSA-based approach only provides a global picture of site conservation for all sequences in the MSA.

Instead of making inferences based on a set of aligned protein sequences, pLMs can take a single sequence as input and make site-specific inferences by considering the amino acid context across all the sequence’s sites. This includes inferring how well each model token (corresponding to the 20 canonical amino acids) fits in every site of the input protein, based on the model’s understanding of protein diversity (inferred using the MLM head layer described above). This pLM property can act as a measure for how likely a specific amino acid substitution is (4, 20). However, we decide to take a different approach at interpreting these probabilities, focusing on assessing how likely a site is to change, instead of trying to determine what amino acid the site is more likely to change to. To achieve this, we calculate the Shannon entropy of the 20 amino acid probabilities per site inferred by the model, into a metric we call “pLM entropy” (Equation 2). This is similar to the alignment entropy described above, with the key difference being that pLM entropies are specific to, and dependent on, each individual input sequence. While a single alignment entropy per site can be calculated for a set of related sequences, every sequence has its own pLM entropy per site, reflecting each specific sequence’s site conservation or variability.

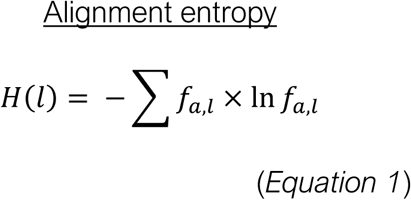

*fa*,*l* being the frequency of amino acid *a* at position *l*.

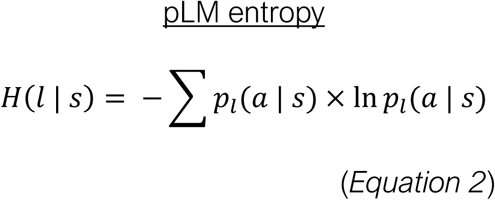

*p_l_* (*a* | *s*) being the pLM-inferred probability of having amino acid *a* given the sequence context *s* at position *l*.

To demonstrate how pLM entropy depends on each sequence’s context, we first calculated pLM entropies for ancestrally inferred internal node sequences in the human H3N2 HA phylogeny. The virus containing this HA protein first jumped into humans during the 1968 H3N2 pandemic and its descendants still circulate in humans until today. In this period of human circulation, the HA protein has exhibited a characteristic ‘ladder-like’ evolution (Figure 2A), providing a clear example for how protein context can change across evolutionary time. We retrieved the internal sequences representing the backbone nodes of the phylogeny, embedded them through our evotuned ESM-2 HA-all model (Figure 1A) and calculated the per-site pLM entropy for every sequence. From the heatmap in Figure 2C, it is apparent that the set of sites that the model considers most likely to change (having higher pLM entropy values) exhibits shifts across the protein’s evolution. These shifts correspond to key protein mutations in the phylogeny taking place over time. For example, site 157 exhibits high pLM entropy between years 2004 and 2006, which is in turn followed by a change from K to I on this site. Similarly, site 162 has high pLM entropy within 2010 and 2011, followed by the site changing from N to S on the tree backbone (Figure 2C). These examples indicate that pLM entropy may hold some predictive signal for which sites are more likely to change in a given sequence context which will be formally explored later in the manuscript.

**Figure 2.**
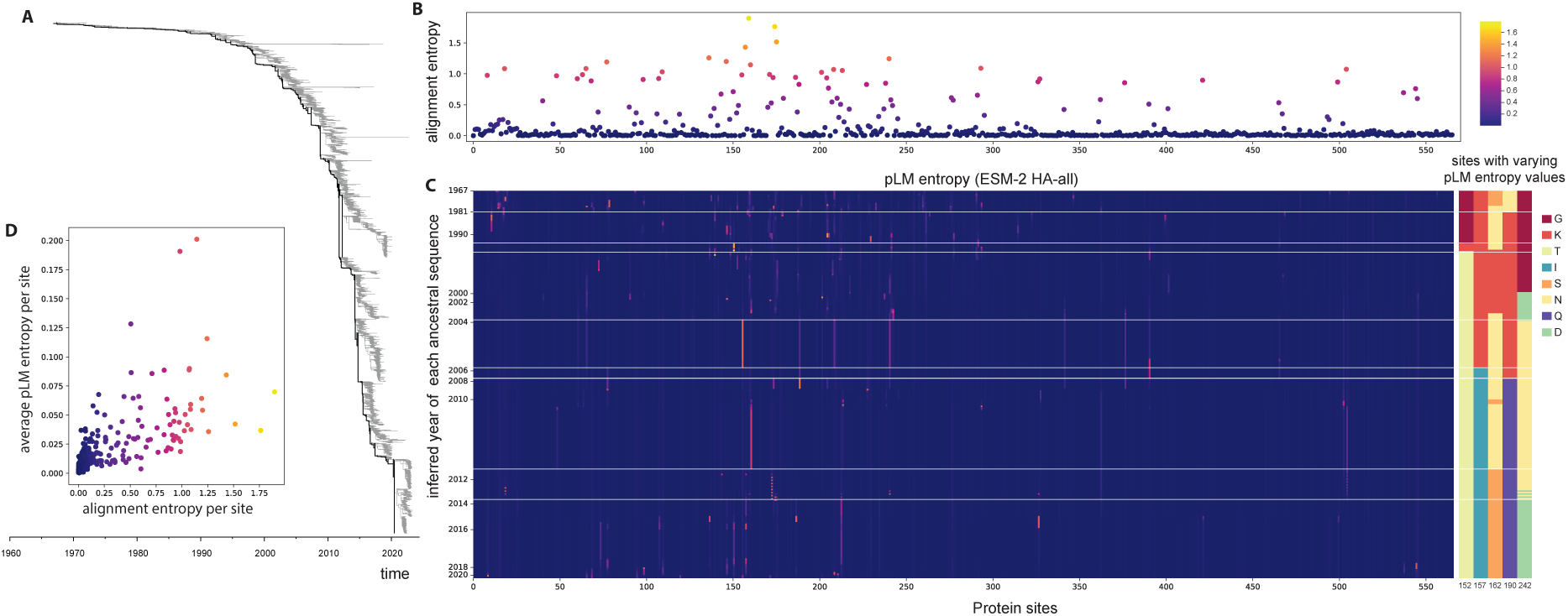
A) Time-calibrated phylogenetic tree of human IAV H3N2 HA sequences. The tree backbone is highlighted in black, corresponding to 265 internal nodes with more than 1000 children nodes each. B) Scatterplot of alignment entropy per site, calculated from the same alignment used for reconstructing the phylogeny in panel A. C) Left: heatmap of per site pLM entropy values inferred for each of the 265 backbone sequences using the ESM-2 HA-all model. Rows are ordered by the node order on the tree. Alignment sites with gaps in any of the ancestral sequences have been removed for clarity. Right: heatmap of amino acid residues present in each backbone sequence for the five sites that exhibited the largest variation in pLM entropy values across the tested sequences. Horizontal white lines indicate notable shifts in the pLM entropy patterns over time. D) Relation between alignment entropy values and average pLM entropy values, corresponding to panel B. Dots are coloured by alignment entropy.

When comparing this pLM entropy pattern to the alignment entropy calculated from all H3N2 tip sequences of the same tree, high and low entropy sites are consistent (Figure 2B). However, our pLM entropy metric allows us to see how this per site residue variation has been shaped across evolutionary time. While average pLM entropy for all the backbone sequences and alignment entropy correlate well (Figure 2D), this global representation of site variation across the tree can be dissected into distinct patterns using the pLM entropy metric on each sequence.

### 3. Evotuned models better capture conservation and variability of HA sites

After demonstrating how pLM entropy relates to alignment-based entropy, we wanted to assess this relationship across all original and evotuned versions of the two models used in this study (Figure 1A). We calculated pLM entropies for sequences from each of the four serotypes used for evotuning (H1, H3, H5, and H7). Unlike the training sequence datasets, the testing sets were not clustered by similarity and contain a larger number of unique sequences (see Methods). We embedded all testing HA sequences through ESM-2, protT5MLM, and all HA-evotuned versions of the two pLMs and calculated their per site pLM entropy. We then constructed MSAs for each serotype and calculated the alignment entropy of amino acid proportions for each alignment column. As described above, pLM entropies are specific to each sequence’s context, while alignment entropies reflect conservation and variability across all aligned sequences (Figure 2). To perform a head-to-head comparison between the two metrics, we first averaged the per site pLM entropies for all sequences in each serotype corresponding to each alignment site (as done for Figure 2C). This resulted in a single average pLM entropy value for every alignment site, reflecting each model’s representation of the global site conservation of each serotype (Figure 1B).

Average pLM entropies per site showed variable levels of correlation to alignment entropies depending on the model used and serotype tested (Figure 3A). Across the comparisons, the HA-all models (both ESM-2 and protT5MLM), evotuned on HAs from all IAV serotypes, achieved the most consistently high correlation (Spearman correlation coefficient between 0.83 and 0.89). The ESM-2 model without evotuning shows moderate correlation but performs substantially better for H1 and H5 (coefficients 0.74 and 0.67) than for H3 and H7 (coefficients 0.43 and 0.45). This is consistent with the phylogenetic relationship between the serotypes (Figure 1B) and likely reflects their relative presence in ESM-2’s original training dataset. Indeed, the UniRef50 cluster contains 51 H1 and 32 H5 HA sequences but 41 H3 and 12 H7 sequences. The protT5MLM model without evotuning showed virtually no correlation in any comparison. This seems to be a result of the base protT5 model not being trained with an MLM-based logit system. However, once all protT5 layers along with the MLM head are evotuned with the HA training datasets, performance becomes directly comparable with that of the ESM-2 evotuned models (Figure 3A).

**Figure 3.**
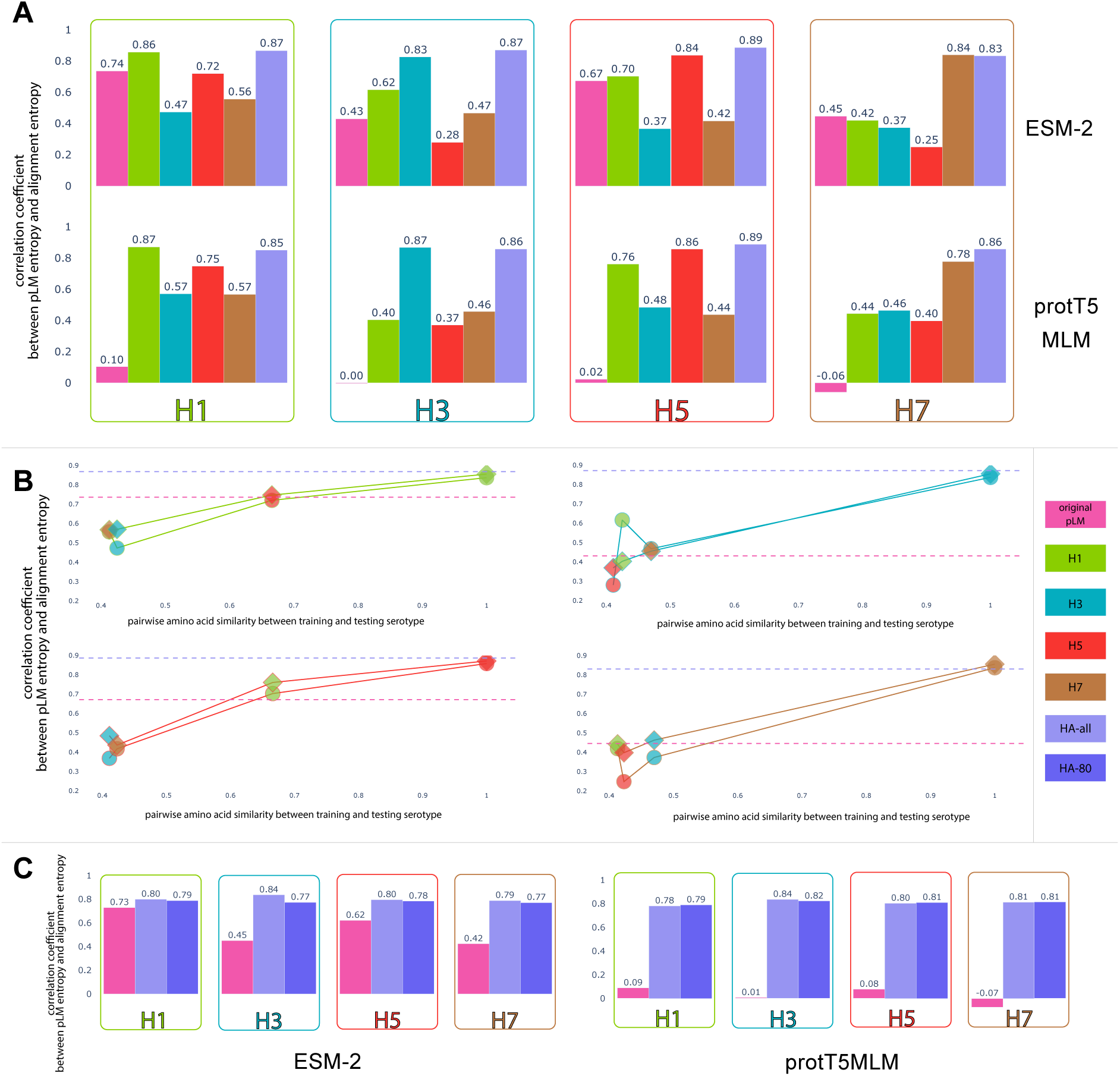
A) Spearman correlation coefficients between average pLM entropy and MSA entropy per site for all four serotypes. ESM-2 models are shown on the top and protT5MLM models on the bottom. Bars are coloured by each model’s training dataset, displayed in the same order as the legend (middle right of the plot). B) Spearman correlation coefficients between average pLM entropy and MSA entropy per site against the pairwise amino acid similarity between testing and training serotype HA sequences. Circles represent ESM-2 models and diamonds protT5 models. Markers are coloured by the training dataset serotype and lines are coloured by the testing dataset serotype (colours corresponding to the legend) C) Spearman correlation coefficients between average pLM entropy and alignment entropy calculated from the 20% most recently collected sequence MSA for the original, HA-all, and HA-80 models.

Models evotuned on single serotypes showed comparable or in some cases higher correlation coefficients than the HA-all models when used on the testing serotype they were trained on. Interestingly, performance was much poorer when these models were applied on a different serotype, reaching coefficients as low as 0.25 (ESM-2 H5 used on the H7 testing sequences). We speculated that phylogenetic distance between the training and testing serotype would explain this trend, so we compared the achieved correlation to the amino acid similarity between serotypes (Figure 3B). We find that for H1 and H5, which share a bit under 70% sequence similarity, using the respective serotype’s model shows comparable or slightly improved performance to the original ESM-2 model. On the contrary, using the models of the much more distant H3 and H7 serotypes (∼40% sequence similarity) on the H1 and H5 testing sequences reduced correlation coefficients well below those achieved by original ESM-2 (Figure 3B). A similar trend is seen for the H3 and H7 testing sets. This suggests that evotuning pLMs on sequences that are substantially genetically distant to the testing sequences (despite still being phylogenetically related) worsens model performance in this site conservation prediction task.

In the above comparisons there is overlap between the training and the testing sequence datasets. Hence, the high correlation coefficients achieved by the evotuned models reflect their ability to learn site conservation patterns by seeing the unaligned sequences instead of the corresponding MSA. We wondered if these patterns stay consistent in the recent evolution of each serotype, potentially giving the pLMs the capacity to extrapolate beyond their training dataset. To test this, we compared the average pLM entropy per site derived from the original models, the HA-all models, and the HA-80 models—trained on the 80% earliest collected HA sequences (before February 2015, Figure 1A)—against the MSA entropy of HAs collected in the latest 20% of the collection date distribution (after February 2015) for each testing serotype. We show that, even though there is no direct overlap between training and testing sequences, the HA-80 models achieve almost identical performance to HA-all, with marginally lower coefficients for the ESM-2 models compared to the protT5MLM ones (Figure 3C).

### 4. pLM entropy can predict sites that changed in a specific sequence context

After validating that pLMs can better represent the global alignment entropy per site once evotuned with phylogenetically related proteins, we decided to explore whether sequence-specific pLM entropy per site holds some unique information about each sequence context and its evolutionary potential. We reconstructed phylogenies for the testing sequences of each serotype, rooted each tree using temporal information and inferred ancestral amino acid sequences for each internal node of the trees along with the substitutions associated with each branch of the tree (Figure 1B). We embedded the ancestral amino acid sequences through the original models, HA-all, and HA-80 models and calculated per site pLM entropies for all of them.

If the pLM entropy metric has some predictive capacity as to which sites are more likely to change in a sequence, then we would expect the entropy for sites that changed in the branch downstream an internal node to be higher than the entropy of sites that did not for that node’s ancestral sequence. We test this by first calculating the difference between each site’s median pLM entropy of all sequences in the tree (internal and external) and each sequence’s pLM entropy value for that site. This gives us a relative representation of each sequence’s placement in the pLM entropy distribution of a given site, where positive values mean that a sequence’s site entropy is in the top 50% of the distribution and negative values in the bottom 50%. We then retrieve all internal node sequences that have had at least one amino acid substituted in the tree and for each substituted site we also randomly pick a non-substituted site from the same sequence. The median normalised pLM entropies of the substituted and non-substituted sites were compared for each tree and the distributions’ similarity was assessed by calculating the overlap between the two distributions’ interquartile ranges (IQRs) (Figure 4).

**Figure 4.**
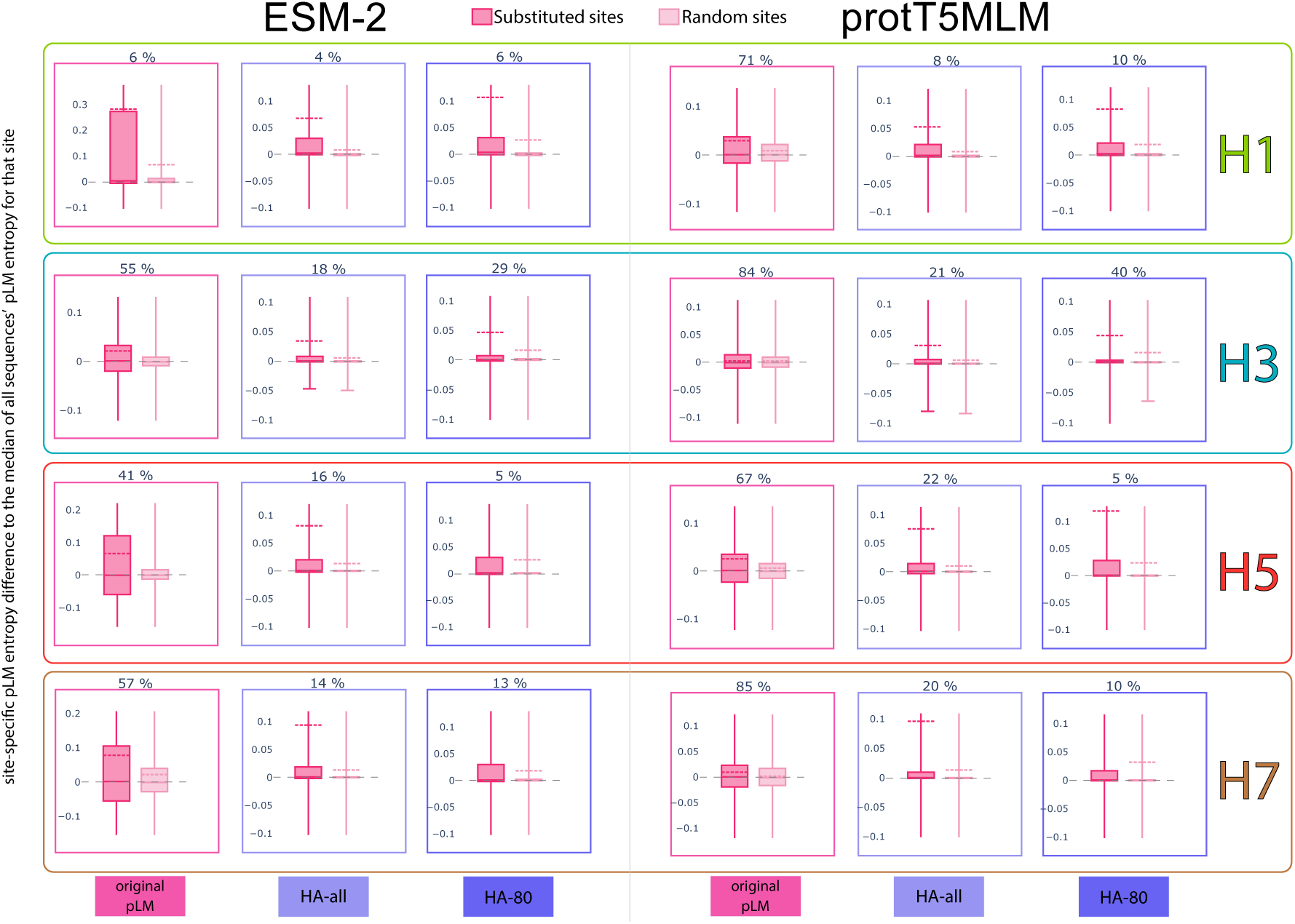
Distributions of the differences between median pLM entropy across each tree and the pLM entropy of internal node sequences that have been substituted for each corresponding site. Values correspond to sites that are substituted in an internal node sequence (hot pink) and an equal number of non-substituted sites from the same sequences (light pink). For clarity, each plot’s y axis range is restricted to 0.1 units above and below the highest 75% IQR and lowest 25% IQR values respectively. Mean values of each distribution are shown as dashed horizontal lines coloured according to the distribution (only shown if they are within the plotted y axis range). The percentage of the substituted site IQR that falls under the 75% IQR point of the random site distribution is annotated on top of each plot. The testing serotype is annotated on the right of each row and the model training dataset is annotated at the bottom of each column (ESM-2 models on the left and protT5MLM models on the right).

For the pLM entropies inferred by the original models (ESM-2 and protT5MLM) the IQRs overlapped almost entirely (Figure 4), suggesting that substituted sites do not have higher entropy values than other sites on the same sequence. The only exception was the original ESM-2’s performance on the H1 testing dataset showing an IQR overlap of 6%, but a very large range for the substituted site IQR. The apparent signal in this model-testing serotype pair is consistent with it achieving the strongest correlation between average pLM entropy and MSA entropy out of the non-evotuned models (Figure 3A). Inferences from the HA-all and HA-80 models (the latter tested only on nodes dated past February 2015) showed much smaller IQR overlap with substituted sites always having higher mean values than the random sites (Figure 4). The majority of substituted site values for the evotuned models were positive, suggesting that the corresponding pLM entropies were in the top 50% of the entropy values for that site. These results imply that, unlike their original versions, the evotuned pLMs infer higher pLM entropy values for sites that are more likely to change in a specific sequence context. This finding is consistent both when sequences from the testing phylogeny are included in the model’s training dataset (HA-all), and when there is no overlap between the sequence datasets (HA-80).

We then decided to assess this apparent signal that sequence-specific pLM entropies demonstrate in a formal predictive framework. For all internal node sequences with at least 3 of the sites substituted on a branch directly downstream that node we fitted a logistic regression model to see if the site-specific pLM entropy for that sequence could predict whether sites would be substituted or not. As a control, we first used the alignment entropy calculated from each serotype’s MSA in the logistic regression. We show that site-specific alignment entropies derived from a single MSA is a poor predictor for whether a site will be substituted or not, successfully predicting (positive slope, p-value < 0.01) between 1 and 3% of the internal nodes tested (Figure 5). On the contrary, using sequence-specific pLM entropies inferred from both HA-all evotuned models showed substantial improvement in predictive power, achieving significant results for up to 12% of the internal nodes tested (ESM-2 HA-all, serotypes H5 and H7, Figure 5). The only pLM that performed worse than the alignment entropy was the protT5MLM model without evotuning, consistent with previous results (Figures 3 and 4). Finally, the percentage of successfully predicted nodes was comparable between the original ESM-2 model and the evotuned models for serotypes H1 and H5. However, the evotuned models still had substantially higher regression slope values than ESM-2, suggesting that, for the nodes with significant regression results, the evotuned models had greater predictive power (Figure 5).

**Figure 5.**
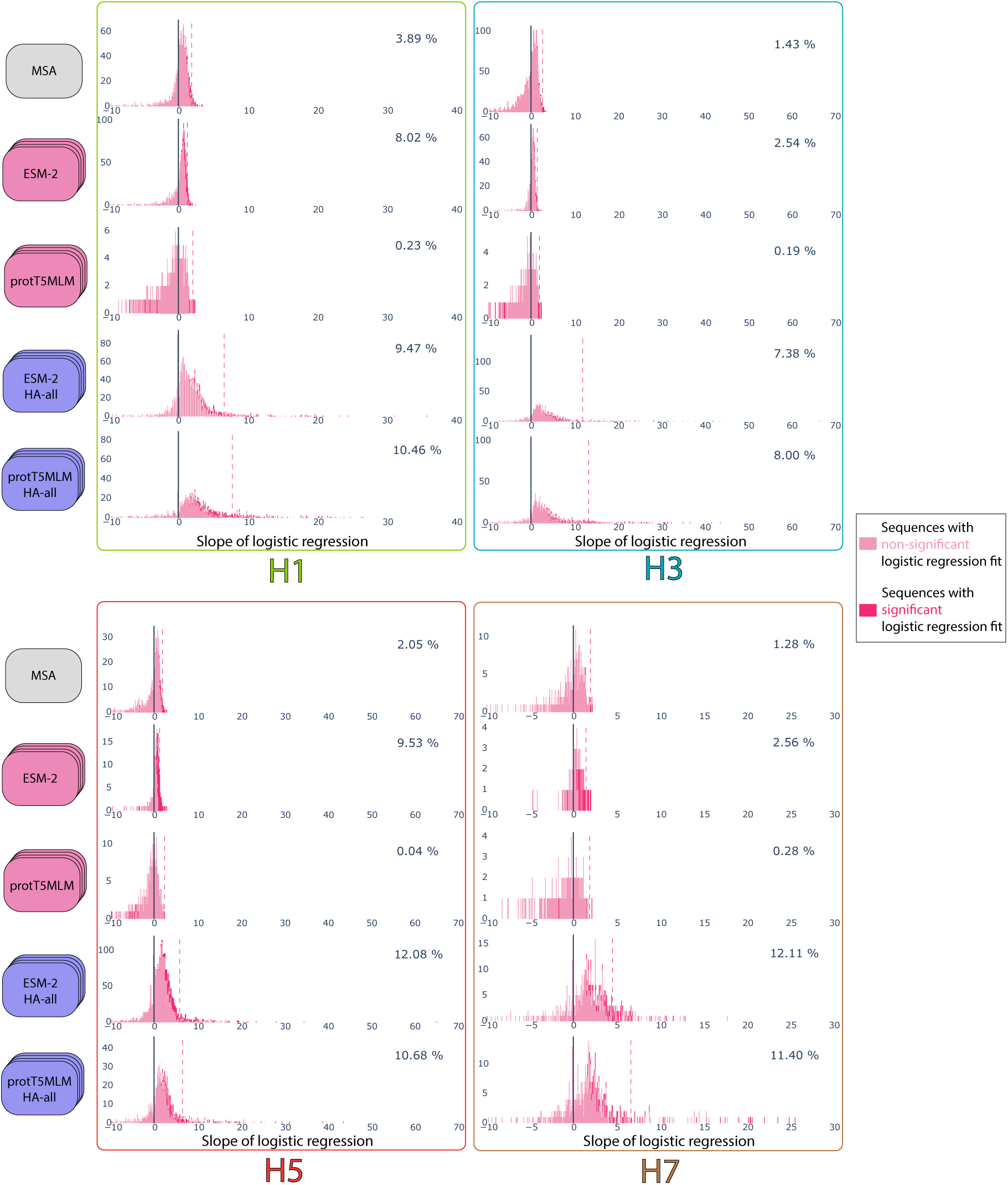
Histograms of slope values for the logistic regression between sites being substituted on an internal node and the alignment entropy (MSA) or the pLM entropy (ESM-2, protT5MLM, and their HA-all versions) of the protein sites. Slopes with values below -10 are not shown in the figures for clarity. Slope values corresponding to regressions with p-values < 0.01 are highlighted in darker pink. The mean slope value for these significant regressions is shown as a dashed line on each histogram. The number displayed on the top right corner of each plot represents the percentage of nodes with significant (p < 0.01), positive slope regressions in each comparison.

Despite the improvement in predictions compared to simply using alignment entropy, pLM entropies still could not correctly predict what sites would be substituted for most of the tested nodes. Rather than proposing that this testing approach presented in Figure 5 can directly be used to predict future evolution, we use this test to showcase that at least some predictive signal about the proteins’ evolution is being learned through pLM evotuning.

## Discussion

If we can create virus-specific pLMs, we can exploit the potential of this novel technology for preparing for future pandemics and better understanding pathogens of concern for public health (35, 36). In this example of applying pLMs to virus proteins we show that evotuning pretrained models on specific sequence groups allows them to better capture the proteins’ diversity and evolution. The pLM entropy metric we propose here is a fitting illustration of how leveraging this new technology enables inferences about conservation and evolutionary potential of specific sites in a protein. Compared to traditional alignment-based entropy per site, our metric provides sequence-specific inferences of site conservation, dependent on each protein’s unique context (Figure 2C). Vigué *et al.* (2022) have previously applied Direct-Coupling Analysis (21) to MSAs to infer context-dependent entropy values for alignment sites (3, 22). Although the authors’ approach follows the same principles as our pLM entropy metric, pLMs do not depend on sequence alignments. Here, we explore the applicability of this alignment-free method for inferring context-dependent protein site entropy.

Our study also provides useful insights on how training sequence availability and their genetic distance to the testing dataset affect pLM inferences in the context of model evotuning. The number of sequences we utilised from each IAV serotype varied substantially in the training datasets, from 4241 H3 sequences to as few as 305 H7 sequences. Still, correlation performance of the H7 evotuned models was directly comparable to models evotuned on the larger datasets (Figure 3A). This is consistent with findings from Biswas *et al*. (2021) showing that model performance was retained even when they reduced their evotuning sequences to 30% of their original dataset (27). Additionally, the HA-80 versions of the two pLMs performed comparatively well with the HA-all versions in both tasks presented here (Figure 3C, Figure 4), indicating that accurate inferences can be made even when there is no direct overlap between the training and testing sequence datasets. We should, however, note that when the training dataset only contained sequences from a different serotype than the one being tested (∼70% sequence similarity) entropy correlation did not improve (at least compared to that achieved by the original ESM-2 model) and when training was done on a distant serotype (∼40% sequence similarity), performance decreased (Figure 3B). Taken together, these results suggest that pLM evotuning is robust to the number of sequences used for fine-tuning, but improving performance depends on how genetically similar the training sequences are to the ones being tested.

The concept of using pLM token probabilities for site-specific inferences, such as mutational fitness and *in silico* deep mutational scanning, has been proposed in a number of recent studies (4, 19, 20). However, most of these approaches focus on predicting the effect of substitutions to a specific amino acid on a given protein site. This is a finer resolution inference which is intuitively harder to make accurately. The pLM entropy metric we describe here is a cumulative measure of these specific substitution probabilities meaning that it is inherently a ‘safer’ inference, while still being biologically relevant. Indeed, in our previous study we showed that aggregating amino acid-specific predictions by *in silico* deep mutational scanning into site-specific scores was an informative approach for downstream analysis (37). Finally, we highlight the potential for pLM entropy, in pair with model evotuning, to be used as a powerful tool to predict the evolutionary trajectory of a protein group (Figure 5).

It should be noted that perplexity is another measure of variation in model prediction, commonly employed in LLMs to evaluate model learning during training. This metric is directly related to Shannon entropy (*perplexity* = *e^H(*s*)^*, where *H*(*s*) is the pLM entropy), but calculated cumulatively across sequences when assessing model prediction. To the best of our knowledge, no previous study has attempted to detect context-specific site variation using per-site perplexity. We favour using entropy instead of perplexity for the purposes of our study so that we can directly compare our results to the alignment-based entropy score, which is the established method for biological sequences.

This study provides a clear demonstration of improving pLM performance through evotuning. From here, there are still many possible adjustments to our methodology that can potentially further advance this relatively new field. For example, we use ancestrally reconstructed sequences only for testing, however in well-sampled trees, confidently inferred ancestral sequences could be used in the training dataset, providing additional unsampled sequence diversity. Including such reconstructed sequences has been shown to improve pLM-based fitness effect prediction (38). The type of evotuning we perform here involves unsupervised learning, focusing on predicting sequence diversity directly captured by the masked learning task. The adaptable nature of pLMs enables other avenues of model training to perform diverse tasks where evotuning remains relevant. The same pLMs fine-tuned through a supervised learning task can predict specific virus phenotypes such as transmissibility (37) or host range (39) from sequence alone. We are confident that evotuning is an important approach for pLM applications, especially when it comes to studying virus proteins which cover an extensive, and still largely undiscovered evolutionary space (40).

## Methods

### Training sequence dataset

HA protein sequences were retrieved from the NCBI Influenza Virus Database(33). All type A sequences, unique on the amino acid level, were downloaded leading to a total of 33657 sequences (retrieved on the 11^th^ of April 2024). Serotype and collection date metadata were also retrieved for all sequences. To remove redundancy the dataset was filtered using MMSeqs2 (41) with a minimum sequence identity threshold of 99% (--cov-mode 0) retaining one representative from each cluster. Sequences with ambiguous characters (X or J) and sequences annotated as being of mixed serotypes were further filtered out leading to a total of 9890 sequences for the final HA-all training dataset. To focus on the four serotypes of greatest concern—H1, H3, H5 and H7—the HA-all dataset was further split by serotype annotation leading to corresponding training datasets of 4241 H3, 2286 H1, 978 H5, and 305 H7 protein sequences.

A final dataset was compiled consisting of the 80% earliest collected HA sequences. We plotted the distribution of collection dates for the 99% filtered HA sequences and determined the 80^th^ percentile point of the distribution to be 2015.16 (February 2015). We then used this cutoff to retrieve a total of 8099 sequences with collection dates before this. Upon manual inspection, we edited the date of isolate A/chicken/Brescia/1902 (ACZ36872) which seems to have an erroneous date recorded from 1902 to 1935 which represents the true collection date (42).

### Model fine-tuning (evotuning)

For fine-tuning the two models we performed an unsupervised masked language learning task (also referred to as domain adaptation) using each IAV HA sequence training dataset described above. For ESM-2, we used the 33 layer, 650M parameter model with the MaskedLM layer (EsmForMaskedLM) downloaded from the Hugging Face library (https://huggingface.co/). Each sequence training dataset was randomly split into an 80% training set and a 20% validation set. The training sequence length was set at 566 amino acids (the length of most HA sequences). Longer sequences were truncated, and shorter sequences were padded to the set length. Consistent with the model’s original training approach, we randomly masked 15% of positions for each input sequence with each masked position having an 80% chance to be a <MASK> token, 10% chance to be a different token, and 10% chance to be the original token. The token type of masked sites was subsequently predicted, and model weights were updated accordingly in batched training steps using a cross-entropy loss function. The training batch size was set at 4 sequences and the validation batch size at 8. The model was trained for a total number of 10 epochs for each training dataset, saving the last 5 epoch checkpoints. The best checkpoint for each fine-tuned model was chosen as the epoch that consistently had both the lowest training and validation loss. Smaller training datasets tended to have an earlier best checkpoint, but all best checkpoints converged between epoch 8 and 10.

For protT5, we used the 24-layer, 1.2 billion parameter protT5-XL encoder-only model. The protT5 model and library do not include a MaskedLM layer, so a custom model, T5EncoderMLM, was designed using the protT5 encoder with an added regression layer module based on EsmForMaskedLM’s EsmLMHead module (https://github.com/huggingface/transformers/blob/6bc0fbcfa7acb6ac4937e7456a76c2f7975fefec/src/transformers/models/esm/modeling_esm.py#L1038). The resulting T5LMHead module takes the last hidden state of the encoder as input and outputs the per-site likelihoods of all amino acids in the model vocabulary. Both the EsmLMHead and T5LMHead modules are constructed sequentially of a linear layer, a gelu operation, a layer normalization, and a final linear layer. Unlike EsmLMHead which has been pretrained with masked sequences, the T5LMHead is initialized with random weights. To minimise the impact of this difference on the comparison of the two models, the T5LMHead weights were pretrained on a set of 2, 909, 523 UniRef50 sequences derived from the UniRef50 IDs used in the ESM-2 pretraining dataset (https://github.com/facebookresearch/esm?tab=readme-ov-file#pre-training-dataset-split--) with sequence length of 1000 residues or less. The protT5 encoder weights were frozen for the pretraining, training only the T5LMHead weights for 2 epochs with a batch size of 8 on the dataset split randomly to 70% training, 20% evaluation, and 10% test subdatasets. Masking methodology and rates were the same as ESM-2 above.

The pretrained T5EncoderMLM achieved a perplexity score of 8.1700. For comparison, ESMforMaskedLM achieved a perplexity score of 6.8178 on the same test dataset. Here, the perplexity score is calculated as *perplexity* = *e*^−∑*_x_ P(x)* ln *Q*(*x*)^ where *P*(*x*) is the true probability distribution and *Q*(*x*) is the probability distribution from the model’s output. Lower values demonstrate higher model certainty in the predictions.

After pretraining, T5EncocderMLM was then fine-tuned on the same datasets with the same hyperparameters on the same hardware as ESMforMaskedLM above. No model layers were frozen for the fine-tuning.

Fine-tuning of both models was done using four NVIDIA RTX 6000 GPUs in Python with Pytorch and the Hugging Face Transformers library (see Data and Code availability).

### Testing sequence dataset

Four testing datasets were retrieved corresponding to the four serotypes also used for model training. The H3 and H1 serotypes primarily circulate in humans and are sufficiently well-represented in the NCBI Genbank database, so the testing sequences for these two serotypes were retrieved from the original dataset from the NCBI Influenza Virus Database described in the ‘Training sequence dataset’ section. The corresponding coding sequences were retrieved from all sequences annotated as being of serotype H1 and H3N2. The full HA and NA serotype was specified for the latter to represent the single clade circulating in humans (which contains the majority of H3 sequences). Coding sequence completeness was ensured by filtering for starting with an ATG start codon, not containing any internal stop codons, and being of length that is a multiple of 3. Sequences that lacked any collection date information were also removed, leading to final datasets of 10151 H3N2 and 8191 H1 coding sequences.

In contrast to the human-circulating serotypes, H5 and H7 viruses primarily circulate in wild birds and are substantially under-represented in diversity in the NCBI Genbank database. Instead, a much larger number of H5 and H7 sequences are available in the GISAID database (https://gisaid.org/). Hence, for the testing datasets of these two serotypes we retrieved all GISAID HA coding sequences annotated as H5 and H7 (accessed on the 27^th^ of June 2024). We filtered out sequences that did not start with a start codon, did not have collection dates or contained internal codons, consistent with the H1 and H3 filtering steps and maintained one sequence for each unique protein translation. This resulted in a total of 8977 H5 and 2229 H7 sequences.

### Sequence alignment and phylogenetic reconstruction

The testing sequence dataset of each of the four serotypes was individually aligned on the amino acid level using the FFT-NS-2 progressive alignment method implemented in mafft v7.525 (43). For aligning the H5 and H7 datasets the gap open penalty (--op) was set to 10 and 7 respectively to avoid misalignment of the indel region containing the polybasic cleavage site insertion common in most high pathogenicity avian IAV HAs. Pal2nal v14 (44) was subsequently used to convert each amino acid alignment to a codon alignment. Each codon alignment was used to infer a phylogenetic tree with IQ-TREE2 v2.3.6 (45) under a GTR substitution model with empirically counted frequencies and accounting for rate heterogeneity with a four category FreeRate model (GTR+F+R4). The resulting maximum likelihood trees were time-calibrated and rooted based on the temporal information with TreeTime v0.11.2 (46). Ancestral sequence reconstruction of protein sequences for all internal nodes and inference of amino acid substitutions along all tree branches was further performed concurrently using TreeTime. Terminal branches of tips with collection dates that did not fit the root-to-tip regression, identified as outliers by the analysis, were manually removed using the ete3 Python3 package v3.1.3 (47).

### pLM entropy and alignment entropy comparison

All HA amino acid sequences and all internal node ancestrally reconstructed sequences for each testing dataset described above were embedded individually (without alignment gaps) through all fine-tuned models as well as the non-fine-tuned 650M parameter ESM-2 model and protT5-XL. The model probability for each amino acid token in each site of an input sequence was retrieved as the softmaxed value from the model’s MLM layer (the existing MaskedLM head for ESM-2 and the added T5LMHead for protT5MLM) (4, 20). Probabilities were then normalised by the highest amino acid probability for the site so that all probability values in a site add up to 1. The pLM entropy per site was calculated as the natural log Shannon entropy of the normalised probability values for each site were calculated using the Python3 scipy package v1.11.1 (48) (Equation 2).

For the testing dataset HA alignment (including only tip sequences), the proportion of each amino acid (excluding gap characters) in each alignment column was calculated. The alignment per site entropy was calculated as the natural log Shannon entropy of amino acid probabilities for each alignment column, similar to the pLM entropy (Equation 1).

Since pLM entropies were calculated for each site of unaligned HA sequences, the resulting values were transformed to match the corresponding alignment positions for each sequence, filling in gap positions with empty values using the pandas Python3 package v2.0.3 (49). The average of per site pLM entropy values of all tip sequences (excluding values of ancestrally reconstructed sequences, consistent with the alignment entropy calculation) corresponding to each alignment column was calculated as the final pLM per site entropy to be compared with the alignment entropy. Sequences with gaps in an alignment column (no pLM entropy calculated) were ignored from the average entropy calculation of the respective alignment column.

The Spearman correlations presented in Figure 3 represent comparisons between each column’s alignment entropy and average pLM entropy. For the ESM-2 HA-80 and protT5 HA-80 model comparisons, only sequences with collection dates falling in the latest 20% tail of the collection date distribution (after February 2015) were considered for the entropy calculations (both alignment and pLM).

### Serotype relatedness analysis

To determine how the HA proteins of different IAV serotypes relate to one another we retrieved the HA sequence with the earliest collection date from each serotype out of our NCBI Influenza Virus Database dataset (serotypes H1 to H18). The reference HA sequence of the Influenza B Virus (NP_056660) was added as an outgroup and these sequences were aligned using mafft (43) (default options) and a phylogeny, presented in Figure 1B, was constructed using IQ-TREE2 (45) under a LG+F+R4 substitution model. The amino acid similarity between serotypes plotted in Figure 3B refers to the pairwise similarity between this set of earliest HA sequences of each H1, H3, H5, and H7 serotypes.

### Comparing pLM entropies between sequence sites

We examined the pLM entropies inferred for ancestrally reconstructed sequences to assess the potential of our metric for predicting sites likely to change in specific sequence contexts.

We first compared sites substituted in the tree to random sites that were not substituted in the same sequence (Figure 4). For each of the four serotype trees we retrieved all internal node sequences corresponding to branches with amino acid substitutions (based on the TreeTime ancestral sequence reconstruction). We then summarised the alignment sites with substitutions in branches immediately downstream these internal nodes. For each of these sites we retrieved the corresponding pLM entropy calculated for the respective internal node sequence by the ESM-2, protT5MLM models and their HA-all and HA-80 versions. We then randomly selected an equal number of sites that did not have any substitutions in the branches directly under each internal node sequence and retrieved their pLM entropies for the same sequence.

Different sites of the protein may be inherently more likely to have higher pLM entropies across input sequence contexts. To remove global, site-specific effects when comparing substituted to non-substituted sites, we normalised their individual pLM entropy values by subtracting the median pLM entropy of the corresponding site distribution for the whole tree (pLM entropies inferred for that alignment sites for all internal and external node sequences, excluding only the internal node being compared). These normalised ‘pLM entropy difference to the median’ values (Figure 4) should be comparable between sites, with positive values indicating that the underling pLM entropy is in the top 50% of all pLM entropies for that site across the tree and negative in the bottom 50% (see Results section 4).

For our second test we fitted logistic regression models to examine if the pLM entropy values had enough context-specific signal to predict the sites that were substituted on an internal node sequence (Figure 5). For all four serotype trees we used the pLM entropies inferred by the ESM-2, protT5MLM models and their HA-all versions. Only values for internal node sequences that had at least 3 sites substituted in any of the branches directly under the node were used for this analysis. For each of these sequences we categorised all sequence sites by whether they had been substituted (1) or not (0) and fit a logistic regression model where substitution category is the dependent variable (y) and per site pLM entropies are the independent variable (X). We used the Logit() function implemented in the statsmodels Python library v0.14.0 (50) and fitted the model using the Broyden-Fletcher-Goldfarb-Shanno (bfgs) algorithm. The slope and p-value of each model was retrieved and visualised in Figure 5.

## Data and Code availability

Code and raw data used for this paper are deposited in the following GitHub repository: https://github.com/spyros-lytras/IAV_HA_pLMs. Model weights are available at: https://doi.org/10.5281/zenodo.14891550.

## Author Contributions

Spyros Lytras conceptualised and designed the study, performed phylogenetic analysis, model evotuning, and model testing, made the figures and wrote the initial draft of the manuscript. Adam Strange developed protT5MLM and performed model evotuning. Jumpei Ito contributed to the study conceptualisation and design. Adam Strange and Jumpei Ito provided editing. Jumpei Ito and Kei Sato contributed to the project administration and funding acquisition. All authors reviewed and proofread the manuscript.

## Acknowledgements

We would like to thank Dr. Michael Heinzinger for useful discussions about the protT5 model. We gratefully acknowledge all data contributors, i.e. the Authors and their Originating laboratories responsible for obtaining the specimens, and their Submitting laboratories for generating the genetic sequence and metadata and sharing via the GISAID Initiative, on which this research is based. This study was partly carried out using the TSUBAME4.0 supercomputer at Institute of Science Tokyo.

